# Evolutionary innovation in conserved regulatory elements across the mammalian tree of life

**DOI:** 10.1101/2024.01.31.578197

**Authors:** Severin Uebbing, Acadia A. Kocher, Marybeth Baumgartner, Yu Ji, Suxia Bai, Xiaojun Xing, Timothy Nottoli, James P. Noonan

**Affiliations:** Department of Genetics, Yale School of Medicine, New Haven CT, USA; Genome Biology and Epigenetics, Institute of Biodynamics and Biocomplexity, Department of Biology, Utrecht University, Utrecht, The Netherlands; Division of Molecular Genetics, Netherlands Cancer Institute, Amsterdam, The Netherlands; Yale Genome Editing Center, Yale School of Medicine, New Haven CT, USA; Department of Ecology and Evolutionary Biology, Yale University, New Haven CT, USA; Department of Neuroscience, Yale School of Medicine, New Haven CT, USA; Wu Tsai Institute, Yale University, New Haven CT, USA

## Abstract

Transcriptional enhancers orchestrate cell type- and time point-specific gene expression programs. Evolution of enhancer sequences can alter target gene expression without causing detrimental misexpression in other contexts. It has long been thought that this modularity allows evolutionary changes in enhancers to escape pleiotropic constraints, which is especially important for evolutionary constrained developmental patterning genes. However, there is still little data supporting this hypothesis. Here we identified signatures of accelerated evolution in conserved enhancer elements across the mammalian phylogeny. We found that pleiotropic genes involved in gene regulatory and developmental processes were enriched for accelerated sequence evolution within their enhancer elements. These genes were associated with an excess number of enhancers compared to other genes, and due to this they exhibit a substantial degree of sequence acceleration over all their enhancers combined. We provide evidence that sequence acceleration is associated with turnover of regulatory function. We studied one acceleration event in depth and found that its sequence evolution led to the emergence of a new enhancer activity domain that may be involved in the evolution of digit reduction in hoofed mammals. Our results provide tangible evidence that enhancer evolution has been a frequent contributor to modifications involving constrained developmental signaling genes in mammals.

## Introduction

Developmentally important genes are often involved in multiple developmental processes, which means they are subject to pleiotropic constraints that impede their evolution. Pleiotropic constraints may be avoided through a modular architecture of gene regulation in which distinct transcriptional enhancers regulate the same gene independently in different developmental time points and cell types [Jeong et al. 2006, Levine 2010]. Evolution at one enhancer may alter the expression pattern of the gene in one context without affecting its expression in other contexts. In this way, the evolution of enhancer sequences can have reduced pleiotropic effects compared to evolution of promoter or protein-coding sequences. Hence, evolutionary modification of the level, timing, and distribution of gene expression may readily occur even for genes that are otherwise strongly constrained [Wray 2007,

Carroll 2008]. Genetic variation in the enhancer sequences of pleiotropic genes has been associated with trait variation in many contexts. For instance, mutations in enhancer sequences have been found to affect traits important in local adaptation [Chan et al. 2009, Wooldridge et al. 2022], and to affect human phenotypes with medical relevance [Benko et al. 2009, De Vas et al. 2023].

Recent studies using biochemical signatures of enhancer function highlighted the frequent loss and rapid evolution of enhancers from non-functional or transposable element sequences [Bourque et al. 2008, Shibata et al. 2012, Villar et al. 2015]. These studies painted a picture of enhancers as weakly conserved sequence elements that turn over at a high rate. While enhancer turnover appears to be a common pattern, many enhancers are evolutionarily conserved over long time scales. In fact, the majority of human *cis*-regulatory elements are highly conserved across mammals [Christmas et al. 2023], and conserved enhancers exert larger regulatory effects than recently evolved ones [Berthelot et al. 2018]. Deeply conserved enhancer sequences have been found to cluster around genes involved in gene regulation and development, pointing to them having a pivotal role in fundamental developmental processes [Woolfe et al. 2005, Visel et al. 2008]. Even deeply conserved enhancer sequences show signs of evolutionary change. For example, Human Accelerated Regions (HARs) are conserved non-coding sequences that show an increase of single nucleotide substitutions specific to the human lineage [Pollard et al. 2006, Prabhakar et al. 2006]. HARs are frequently located near genes involved in neurodevelopment, indicating they may play a role in the development of human-specific adaptations related to brain evolution [Pollard et al. 2006, Prabhakar et al. 2006]. Experimental interrogation has shown that HARs encode human-specific enhancer functions, particularly related to neurodevelopment [Prabhakar et al. 2008, Boyd et al. 2015, Doan et al. 2016, Geller et al. 2019, Won et al 2019, Aldea et al. 2021, Girskis et al. 2021, Uebbing et al. 2021, Dutrow et al. 2022, Whalen et al. 2023]. Hence, the evolution of conserved enhancers may have contributed to important developmental adaptations across mammals by enabling evolutionary change in conserved, pleiotropic genes.

To better understand the extent and the effects of evolutionary innovation in conserved enhancer elements, we studied signs of evolutionary sequence change in conserved, putative enhancer sequences throughout the mammalian tree of life. Mammals are an extraordinarily diverse group of organisms that live in many different environments, such as underground, on land, up in trees, under water, or in the air. Such vastly different lifestyles require physical adaptations that evolved, sometimes more than once, in many independent mammalian groups [Luo 2007]. This suggests that the developmental processes and pathways underlying these adaptations were modified repeatedly during mammalian evolution. In addition, the genetic diversity of mammals has been sampled extensively and whole genome sequences are available for hundreds of mammalian species [Hecker & Hiller 2020, Kirilenko et al. 2023]. To study the evolution of conserved enhancers across mammals, we collected a set of putative regulatory elements conserved within mammals and tested for acceleration of sequence substitutions (below shortened to “(sequence) acceleration”) within these elements along internal branches of the mammalian tree of life. We found evidence of pervasive acceleration throughout the mammalian phylogeny. We also found that the substitutions underlying the sequence acceleration signal potentially impact enhancer function by frequently altering transcription factor binding sites. Regulatory elements of genes with roles in the immune system were more frequently affected by sequence acceleration. However, genes with a complex gene regulatory architecture experienced more acceleration over their many regulatory elements combined, even more than immune system genes. These include deeply conserved and pleiotropic developmental transcription factors such as members of the Notch signaling pathway. This supports a model in which gene regulatory evolution facilitates evolutionary change affecting deeply conserved, pleiotropic developmental pathways.

## Results

### Accelerated evolution in putative enhancers is pervasive across the genome and the mammalian tree of life

To study sequence acceleration in enhancer elements genome-wide across the mammalian phylogeny, we used a recently published whole genome alignment containing the genomes of 120 mammal species (Fig. 1A) [Hecker & Hiller 2020]. As our candidate enhancer elements we collected two sets of putative regulatory elements (PREs) that are conserved across most of the whole genome alignment dataset. First, we used phastCons elements (pCEs), which are defined as constrained elements identified using the 120-mammal whole genome alignment [Siepel et al. 2005, Hecker & Hiller 2020]. These sequences contain some of the oldest and most deeply conserved enhancer sequences. It was previously shown that pCEs are enriched in regulatory elements [Visel et al. 2008, Holloway et al. 2016]. Second, we used human candidate *cis*-regulatory elements (cCREs) identified by the ENCODE consortium [ENCODE et al. 2020]. We reasoned that a certain degree of conservation is needed to effectively test for acceleration across diverse mammals. We therefore selected elements that also showed biochemical signatures associated with active chromatin in mouse (Methods) to remove weakly conserved elements. ENCODE cCREs were defined as regions with high prevalence of biochemical signatures of enhancer activity (Methods) [ENCODE et al. 2020]. The majority of human cCREs are conserved across mammals [Christmas et al. 2023]. By selecting sequences whose activity is conserved between human and mouse, we further enriched for evolutionarily conserved cCREs, although not typically to the degree of pCEs (mean phastCons, pCEs, 0.761 vs. cCREs, 0.299, Mann-Whitney *U* = 1.6 · 10^9^, *P* < 10^-307^; Fig. S1; [Siepel et al. 2005]). cCREs were still more highly constrained than the genome average (mean phastCons over the entire non-coding genome, 0.101; Wilcoxon *T* = 5.9 · 10^9^, *P* < 10^-307^; Fig. S1). This increased level of conservation was also reflected in the fact that ENCODE cCREs significantly overlapped pCEs (43.8% of all cCREs overlapped a pCE; resampling *P* < 0.0001). We filtered these two sets of sequences against exons, promoters, pseudogenes, and repeats. The resulting dataset of putative regulatory elements contained 319,292 pCEs and 115,014 ENCODE cCREs for a total of 434,306 PREs covering 5.7% of the non-coding, non-repetitive human genome. We defined these PREs in the human genome, and they map to orthologous sequences in other genomes throughout the alignment. We note that the number of elements is high due to the fact that many PREs may represent segments of larger enhancer sequences defined in other ways.

**Fig. 1.**
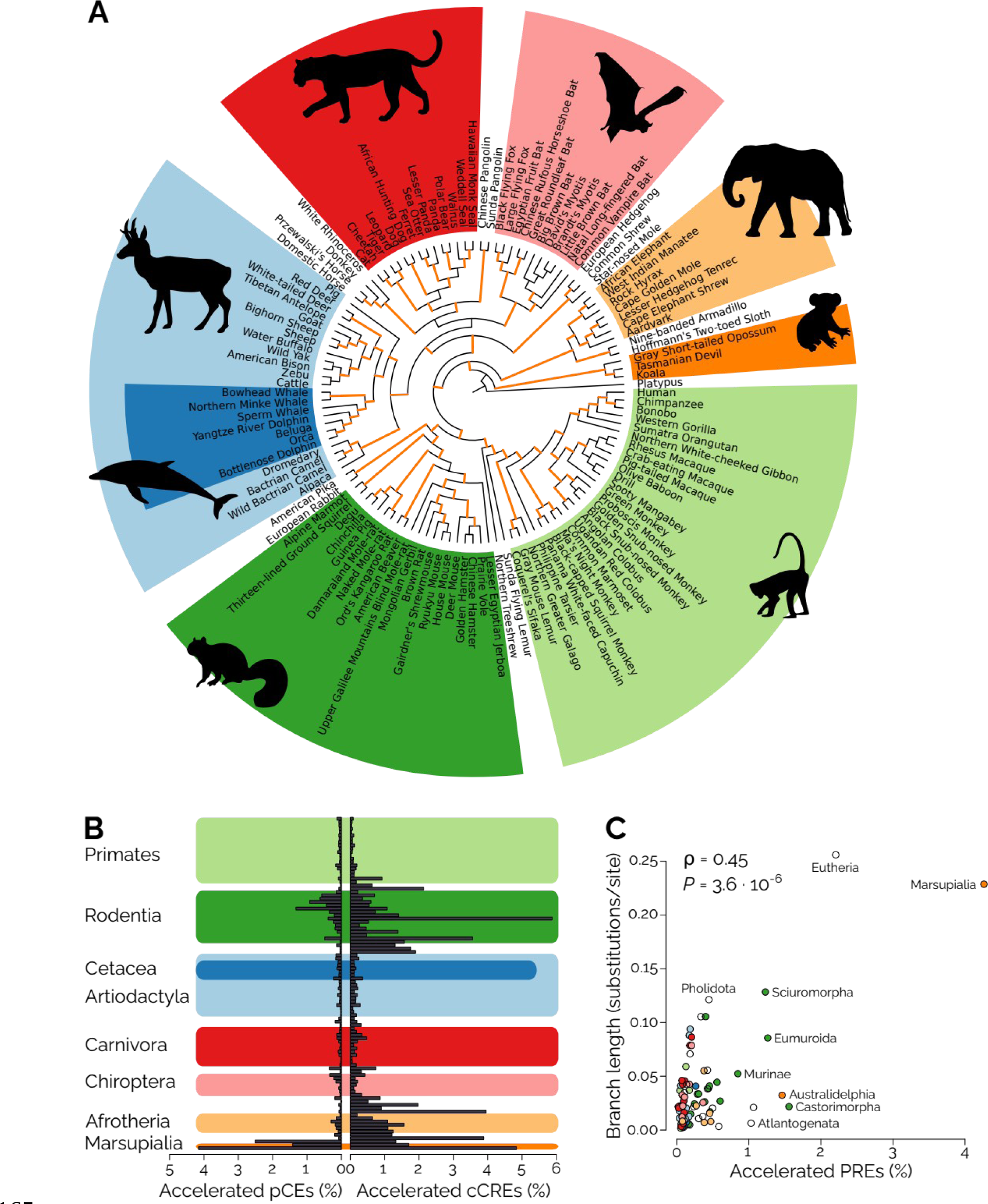
Overview of the dataset used to identify accelerated putative regulatory elements. (**A**) A tree of life showing the relationships between the 120 mammal species in the alignment. PREs were tested for acceleration on 100 internal branches (orange). Colored blocks highlight major mammalian clades, labeled in (B). (**B**) Fractions of pCEs and cCREs found to be accelerated on the 100 tested branches. See Fig. S3 for an expanded version with all branch names. (**C**) Branch length correlates with the proportion of PREs exhibiting acceleration. Colors as in (A) and (B).

To identify PREs that exhibit evidence of sequence acceleration, we used phyloP [Siepel et al. 2006] and tested for acceleration of sequence substitutions on 100 selected internal branches [Holloway et al. 2016] of the tree spanning the 120 mammalian species (Fig. 1A; Methods). We focused on internal branches, rather than terminal branches, because comparing single genomes to the rest of the alignment can be difficult to interpret. In particular, the genetic variation that distinguishes a single genome sequence from the other genomes in the alignment can represent two cases: 1, variants that are polymorphic within the species that is represented by the single genome sequence, and 2, variants that are fixed within the species. When using only one genome sequence representing each species, it is impossible to know which variant relates to which of the two cases. We therefore restricted our analysis to internal branches which divide the alignment dataset into two groups of genomes, but not single genomes. Variants that distinguish the two groups on either side of an internal branch are likely to be fixed in those two groups and thus the type of variant that would underlie lineage-specific adaptations. We also removed any species’ PRE sequences if they contained a large amount of insertions or deletions (>120% or <80% of the human PRE length; Methods); and we removed any test results that were likely affected by GC-biased gene conversion (Methods), a substitution bias that can lead to an increased amount of substitutions in the absence of positive selection. These filters removed individual PRE-by-branch combinations and not entire PREs, but 66 PREs were removed in every instance and hence altogether from the dataset; the remaining 434,240 PREs were only removed at a few branches each but not across the whole tree (Fig. S2).

We applied phyloP to test acceleration in these PREs on every branch ortholog that was not filtered out, for a total of 37,394,226 tests. Overall, 108,737 tests (0.29%) were significant after multiple-testing correction (Fig. 1B, S3). 81,195 PREs (18.7%) were accelerated on at least one branch. The vast majority of PREs were accelerated on a small number of branches tested: The mean number of branches on which a PRE showed evidence of acceleration was 1.3. The Marsupialia branch (separating marsupials from the rest of the tree) showed the largest number and proportion of accelerated PREs (9,389, 4.26%). As is expected from the fact that ENCODE cCREs have lower conservation levels than pCEs, cCREs were more frequently accelerated than pCEs (0.59% vs. 0.20%). cCREs also showed higher proportions of acceleration than pCEs on 86 out of the 100 branches tested for acceleration and the highest overall frequency of acceleration was found among ENCODE cCREs accelerated in Castorimorpha (the branch uniting beavers and kangaroo rats; 5.9%, Fig. 1B, S3). Longer branches showed more sequence acceleration (correlation between branch length and frequency of acceleration of all PREs tested in a branch, Spearman’s ρ = 0.45, *P* = 3.6 · 10^-6^, Fig. 1C), indicating that the overall degree of divergence among lineages is an important factor in explaining the amount of sequence divergence in the form of acceleration we observe.

To gain insight into the genes that may be regulated by our set of PREs, we collected genome-wide chromatin conformation capture-type datasets generated from human and mouse developmental samples (including developing tissues and developmental model cell types) to assign PREs to putative target genes (Methods). We also assigned PREs residing in close proximity to a promoter (1–5 kB upstream of a transcription start site) to that gene. We connected 195,123 PREs (44.9%; 74,364 ENCODE cCREs and 120,759 pCEs) to 39,323 unique ENSEMBL genes. Most connections (83.2%) were supported by chromatin conformation data; the remainder were supported by promoter proximity (15.1%) or both types of connections (1.71%). PREs were connected to an average of 3.02 genes. ENCODE cCREs were on average connected to slightly more genes than pCEs (3.57 vs. 2.68, respectively). Genes were connected to an average of 15.0 PREs (range 1–798), a value inflated by the fact that many PREs may represent subparts of larger enhancers. Having established connections between PREs and genes, we could test for enrichment of Gene Ontology (GO) categories and found that both ENCODE cCREs and pCEs were connected to genes involved in diverse processes (Tab. S1); particularly pCEs were connected to genes involved in gene regulation and development, as has been observed before [Woolfe et al. 2005, Pennacchio et al. 2006].

### Degrees of evolutionary constraint affect levels of acceleration

We first sought to identify factors that influence which PREs showed evidence of acceleration. We examined sequence constraint, as PREs that are subject to strong purifying selection are likely to be less tolerant of sequence acceleration. Sequence constraint as measured by phastCons over the length of a PRE correlated negatively with its frequency of acceleration (Spearman’s ρ = -0.14, *P* < 10^-307^; Fig. 2A), indicating that more constrained PREs indeed show less acceleration.

**Fig. 2.**
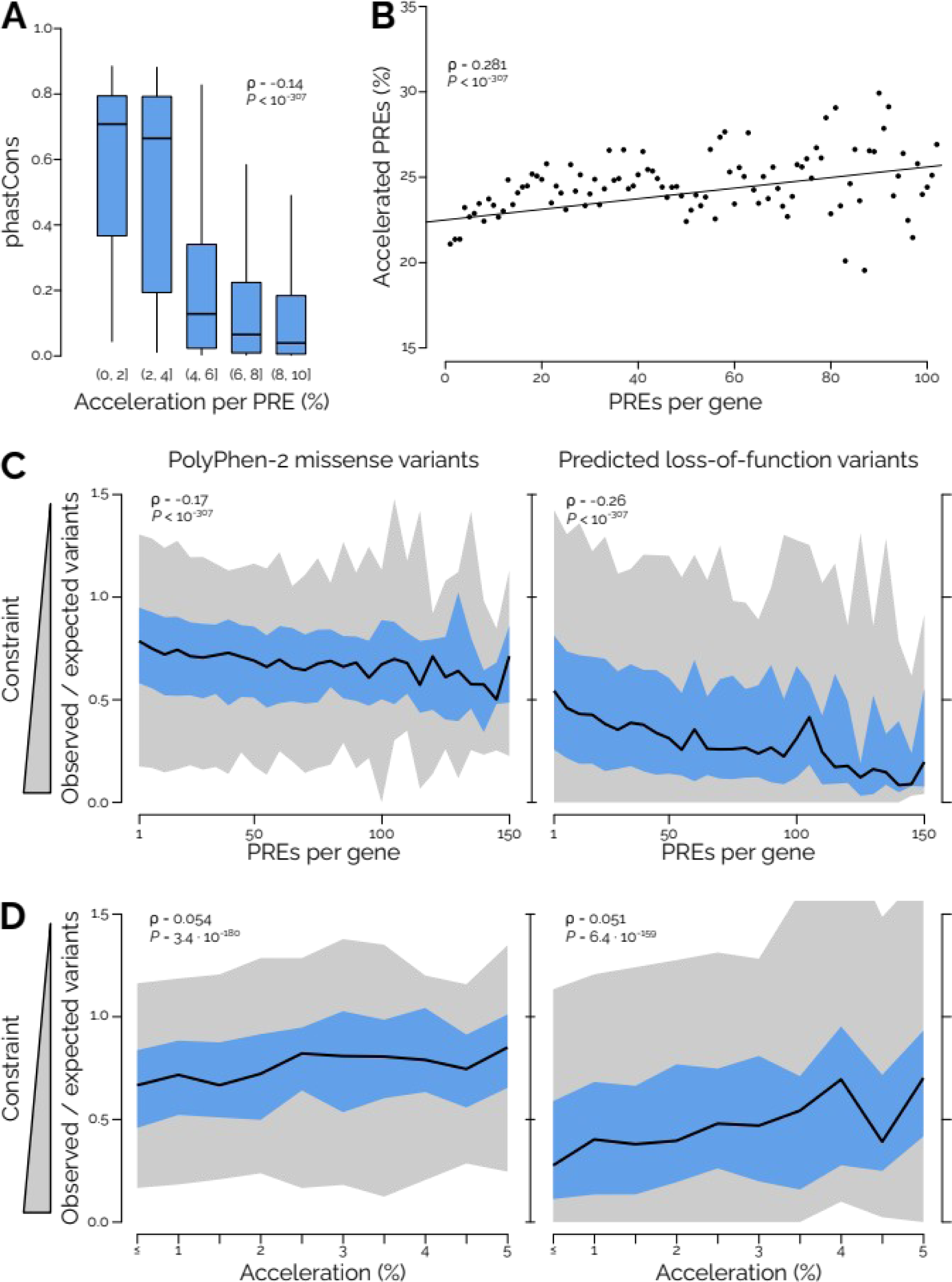
Constraints associated with the level of acceleration in PREs. (**A**) Sequence constraint inferred by phastCons averaged over the length of a PRE is correlated negatively with its acceleration frequency. PREs were divided into five groups according to how frequently we found them accelerated (percent accelerated among branches tested). Boxes show the inter-quartile range of phastCons for each group, with median indicated by a bar; whiskers, 90% data interval. Only PREs with <10% acceleration are shown because data above this value was sparse. (**B**) Genes with fewer PREs show less acceleration in their associated PREs. We found an overall effect of the number of PREs per gene on how frequent those PREs were accelerated. For visualization purposes, dots show the mean number of PREs that are accelerated at least once per gene, averaged over all genes with that number of PREs but correlation testing was performed on the raw data (how many PREs per gene were accelerated on at least one branch vs PREs per gene). (**C**) Genes that are highly constrained according to gnomAD data tended to be in contact with larger numbers of PREs. The y axis shows inactivating variant tolerance from gnomAD as the observed/expected frequency of missense or loss-of-function variants per gene. The x axis shows number of PREs per gene. We found negative correlations between variant tolerance and the number of PREs per gene. Data was analyzed in PREs per gene-bins of size 5; dark line, median; blue area, inter-quartile range; gray area, 95% data interval. (**D**) Genes that are highly constrained according to gnomAD data tended to show less acceleration in their associated PREs. The x axis shows acceleration frequency, measured as the number of significant acceleration tests relative to all the acceleration tests for all PREs of a gene. We found positive correlations between variant tolerance and acceleration frequency. Colors as in (C).

We next examined whether the number of regulatory elements associated with a gene influences the propensity of these regulatory elements to undergo sequence acceleration. One might hypothesize that genes with very few regulatory elements would show increased constraint within those regulatory elements, because those few elements contain all of the information that regulates those genes. Conversely, genes with particularly large numbers of regulatory elements may show less constraint in those elements, as the regulatory information regulating the gene is distributed over a large sequence space and any one element may contain relatively little information. Therefore, we investigated whether the proportion of accelerated PREs is affected by total PRE number for a given gene. We found a slight overall correlation between the number of PREs per gene and the proportion of accelerated PREs per gene (Spearman’s ρ = 0.281, *P* < 10^-307^). This suggests that for genes with few regulatory elements, those elements are under higher sequence constraint than those of genes that have many regulatory elements.

Another possible source of constraint may come from the constraints on the gene itself. For example, more constrained genes could have more constrained regulatory elements. We tested if genes under strong constraint also show a lower proportion of accelerated PREs. We cross-referenced the genes contacted by our PREs with the gnomAD constraint database (v. 2.1.1; [Karczewski et al. 2020]), which classifies genes according to their intolerance for inactivating mutations. We focused on two measures of mutation intolerance, the prevalence of predicted loss-of-function mutations and the prevalence of PolyPhen-2 missense mutations, which is a method predicting the impact of missense mutations on protein function [Adzhubei et al. 2010]. We indeed found correlations between target gene constraint and acceleration in PREs, although the amount of variance explained was small. First, we found that genes with lower frequencies of PolyPhen-2 missense variants or loss-of-function variants tended to have proportionately more PREs (Spearman’s ρ = 0.17–0.26, *P* < 10^-307^; Fig. 2C). Second, we found a lower proportion of accelerated PREs associated with genes with fewer deleterious variants (Spearman’s ρ = -0.051 – -0.054, *P* < 10^-158^; Fig. 2D). These two results suggest that more constrained genes 1) are associated with a more complex *cis*-regulatory architecture, and 2) show less evolutionary change within their regulatory elements.

### Sequence acceleration correlates with putatively functional changes in PREs

Next we explored whether acceleration in PREs pointed to sequence changes that potentially altered their function. We asked whether sequence acceleration correlates with increased transcription factor binding site (TFBS) turnover. To do so, we generated genome-wide TFBS predictions for all 120 genomes in the alignment. We then considered each branch at a time, and compared the sequence content of the group of species united under this branch with that of the species in the immediate outgroup (Fig. 3A). We required branches to have more than one species in their immediate outgroup (the ingroup always fulfilled this criterion because we were testing internal branches only). For instance, the branch leading to genus *Pan* (chimpanzee and bonobo) has only human in its immediate outgroup and was excluded. For each branch and PRE, we then measured how dissimilar the TFBS content was between the ingroup species and the outgroup species (Methods). Once we had calculated dissimilarity values for every PRE-by-branch combination, we compared all cases in which PRE-by- branch combinations were accelerated with those where we did not detect acceleration (Fig. 3A). We found that significant acceleration coincided with increased rates of TFBS turnover: The dissimilarity between the TFBS sets between ingroup and outgroup was larger in cases where the PRE on that branch was accelerated (over the entire dataset, median dissimilarity between ingroup and outgroup when accelerated, 0.85, vs. when not accelerated, 0.79; Mann-Whitney *U* = 1.3 · 10^11^, *P* < 10^-307^; Fig. 3B). When tested in each branch individually, 55/59 branches showed a significant difference between accelerated and non-accelerated PREs and all of those 55 branches showed a larger dissimilarity when accelerated. This supports that sequence acceleration frequently altered TFBS content, consistent with natural selection for altered function as a driving force behind accelerated sequence divergence within regulatory elements.

**Fig. 3.**
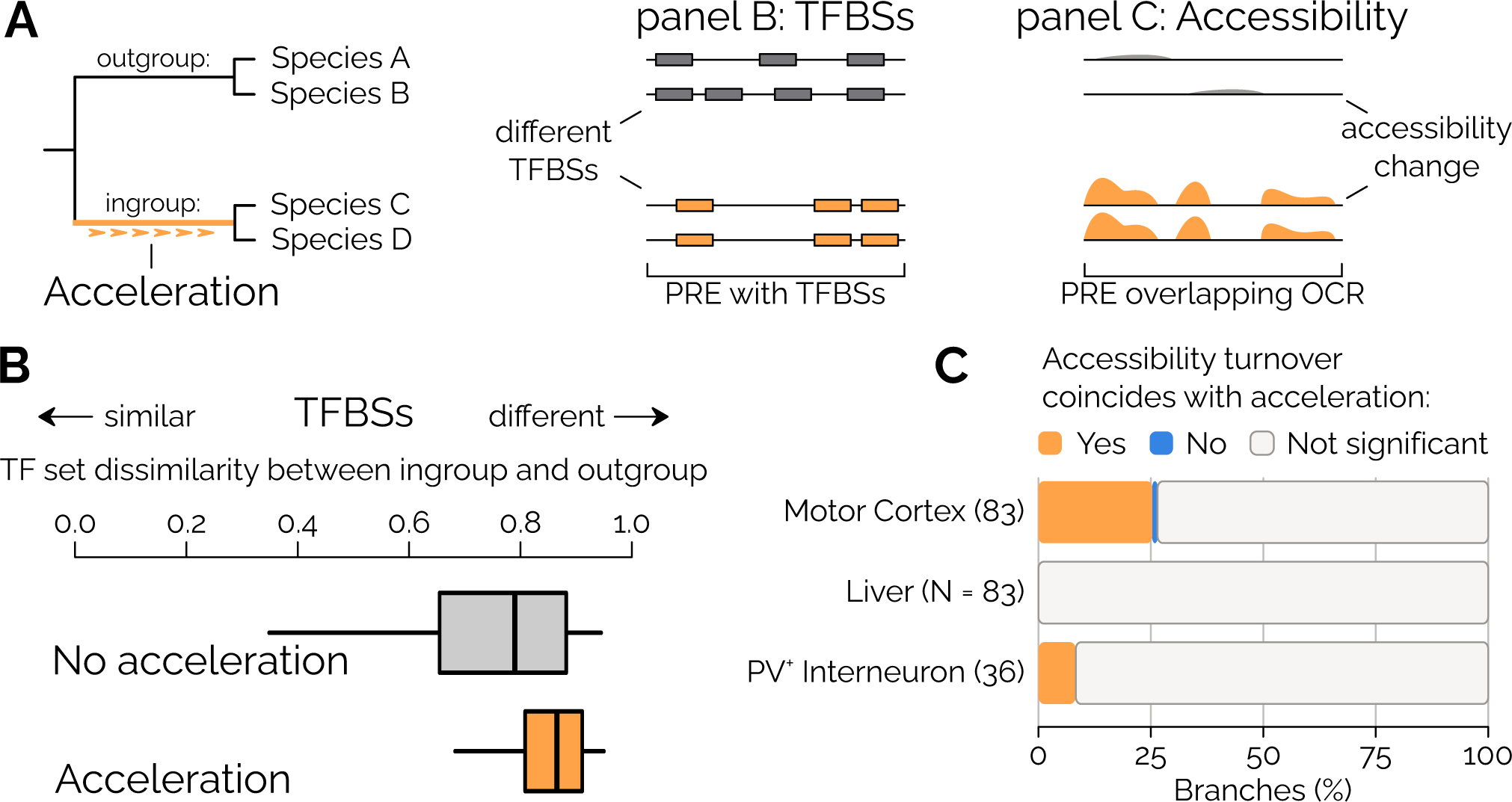
TFBS turnover is increased at accelerated sequences. (**A**) Schematic illustrating our analytical approach. Left: An acceleration event on a branch (uniting species C and D) leads to sequence divergence of this clade relative to the outgroup (species A and B). Middle: Turnover of predicted TFBSs in a PRE coincides with the acceleration event. Right: Turnover of chromatin accessibility in a PRE coincides with the acceleration event. (**B**) The sets of predicted TFBSs in ingroup and outgroup in the whole dataset are more different in cases where the sequence is accelerated. Box plot show the TFBS set dissimilarity between ingroup and outgroup for instances where there was no significant acceleration vs. instances were we detected significant acceleration. The distributions of dissimilarity estimates over all PRE-by-branch combinations divided into those two cases are shown by boxes of inter-quartile range with median indicated by line; whiskers, 90% data interval. (**C**) Accessibility turnover in open chromatin regions (OCRs) often coincides with sequence acceleration. We compared OCR accessibility predicted using machine learning (Kaplow et al. 2023) with acceleration in our PREs and found that there is often a larger difference in predicted accessibility change between ingroup and outgroup when the PRE was accelerated on that branch. We could intersect 83 branches each with motor cortex and liver data and 36 branches with interneuron data. The plot shows the proportion of branches in which we found statistically significant accessibility change between ingroup and outgroup for each OCR dataset. Orange denotes an openness change that coincides with acceleration on that branch while blue denotes a significantly smaller change in openness on an accelerated branch. Most branches did not show significant differences in accessibility change between in- and outgroup for accelerated versus non-accelerated PREs (gray). When there was a significant difference, acceleration coincided with OCR turnover in all but one case (motor cortex tested on branch Pholidota, pangolins).

In a similar approach to comparing TFBS turnover to acceleration, we compared our acceleration results to a recent study that used machine learning to infer the activity of gene regulatory elements across mammals (Fig. 3A) [Kaplow et al. 2023]. This study identified open chromatin regions (OCRs) using ATAC-seq in primary samples from liver, motor cortex, and parvalbumin-positive (PV^+^) interneurons. The authors then used OCR data from just a few species (2–5; Methods) to infer the chromatin state in each of the three sample types in orthologous sequences of up to 222 mammal species using machine learning based solely on genetic variation found in those species’ orthologs. Given that our two approaches both use genetic variation to identify patterns of evolutionary change, we sought to determine whether the two approaches identified a similar set of regions as undergoing evolutionary change by comparing sequence acceleration with changes in accessibility. We considered all OCRs that overlapped a tested PRE. For each OCR and for each branch, we calculated the average openness among ingroup and among outgroup species and took the absolute difference between them. We then compared these differences between all OCRs where the branch was accelerated versus non-accelerated. In the example sketch (Fig. 3A), we would calculate the average openness in each OCR in ingroup and outgroup (Fig. 3A) and form the absolute openness difference between these two groups for each OCR. Then we compare these openness differences between all OCRs overlapping accelerated PREs with all OCRs that were not accelerated. Our expectation was that we would observe a larger difference in openness between in- and outgroup for OCRs overlapping accelerated PREs than for OCRs overlapping non-accelerated PREs, if acceleration was associated with change in enhancer activity. And indeed, 22 of 23 branches with a significant openness difference between ingroup and outgroup in the motor cortex dataset showed the larger openness difference when the overlapping PREs were accelerated on that branch than when the PREs were not accelerated (Fig. 3C). In the PV ^+^ interneuron dataset, all three branches with a significant difference openness difference showed the larger openness difference at accelerated PREs. In the liver dataset, none of the branches showed a significant openness difference between ingroup and outgroup. In the other two datasets most branches showed also no significant difference in openness between accelerated branches and non-accelerated branches (Fig. 3C). One reason for the large number of insignificant test results may be that the dataset we could analyze was small, because 1) acceleration is rather rare on many branches, and 2) the set of intersected sequences of PREs and OCRs is small, as these two sequence sets each encompass small portions of the genome. The number of OCRs overlapping accelerated PREs per branch was largest in the motor cortex dataset (mean, 71.7) but much smaller in liver (28.3) and in PV^+^ interneurons (21.1). In summary, while we may lack sufficient data to see a stronger pattern, we found an alignment between the acceleration signal in our study and turnover of chromatin accessibility predictions [Kaplow et al. 2023].

### Genes with developmental and gene regulatory functions have many accelerated PREs

We set out to explore the biology that is putatively affected by sequence evolution in our PREs through gene set enrichment analyses in a series of questions. First, we used GO term enrichment testing to ask which biological processes were enriched among genes with accelerated regulatory elements on each branch separately. We identified GO terms enriched in genes with accelerated PREs on a given branch compared to all genes associated with a PRE. Of the 100 analyzed branches, 85 showed significant enrichment for GO categories (Tab. S2). Different branches showed enrichment for different types of GO categories. We adapted a GO slim approach for complexity reduction (Methods) and found that GO categories that were repeatedly enriched on multiple branches included GO terms related to the broad categories of cell differentiation, development, immune system functions, and response to stimulus (Fig. S4). In summary, branches showed enrichment for different GO categories, but some types of GO categories were more frequently enriched on different branches than others.

Because our analysis covers branches across the entire mammalian phylogeny, we were able to assess how often PREs are accelerated on more than one branch, referred to here as “repeated PRE acceleration.” Of all accelerated PREs, we found 24.8% (20,152) accelerated on more than one branch, most of which (15,143) were accelerated on two branches and the rest on more than two branches. To gain insight into the biology of repeated PRE acceleration, we characterized PREs through their associated genes. First, we calculated the average acceleration frequency of all PREs contacting the same gene. We then took the top 5% of genes with the highest average PRE acceleration frequency and tested for GO and Reactome enrichment, relative to all genes connected to PREs. We chose a 5% cutoff because it provided large enough gene numbers for effective enrichment testing. We found that these genes were enriched for biological processes related to immune response (Tab. S3), suggesting that adaptive evolution involving gene regulatory elements is most frequently due to pathogen defense processes.

Next, we reasoned that the genes with the largest numbers of PREs would have the greatest potential for repeated evolution via sequence acceleration in PREs. Similar to above, we compared the top 5% of genes connected to the largest numbers of PREs to all genes connected to PREs and performed GO and Reactome enrichment analysis. Many of the enriched terms we found fall into two broad categories of biological processes, development and gene regulation (Tab. S4). Examples of the most highly enriched terms include *regulation of transcription by RNA polymerase II*, *anterior/posterior pattern specification*, *proximal/distal pattern formation*, and *nervous system development* (Tab. S4).

Finally, we asked which kinds of genes showed the most acceleration over all their PREs combined. A gene that is contacted by many PREs may experience a lot of acceleration over all of its PREs, even if those individual PREs are not as frequently accelerated as others. In fact, each gene that is contacted by at least one accelerated PRE had accelerated PREs on 5.5 different branches on average. When we tested for GO and Reactome enrichment among the top 5% of genes with the highest total number of acceleration events over all of their PREs combined, we again saw an enrichment for biological processes related to gene regulation and development, but not for immune response (Fig. 4A, Tab. S5). This result suggests that genes that experience frequent evolutionary innovation in their gene regulatory elements tend to be involved in gene regulation and development. However, these genes experience frequent evolutionary innovation because they possess large numbers of regulatory elements rather than elements that show high acceleration frequencies individually. As we showed above, PREs with high individual acceleration frequencies are contacting genes enriched in immune system functions but not gene regulation or development.

**Fig. 4.**
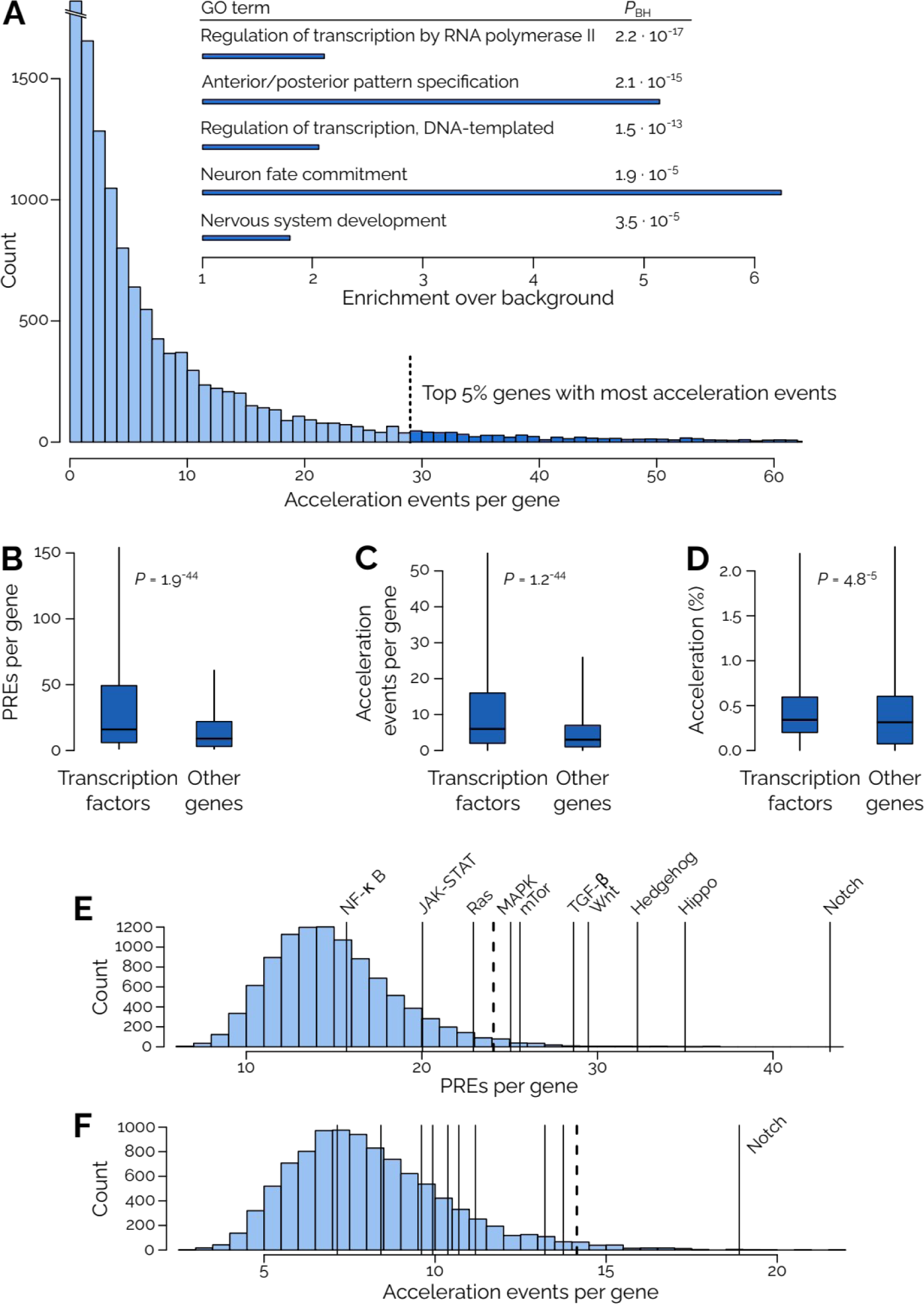
**Genes involved in gene regulation and development are enriched in accelerated PREs**. (**A**) The 5% of genes with the most acceleration events are enriched for GO terms related to gene regulation and development. Examples of highly enriched GO terms among the 5% genes with most acceleration events. For a full list see Tab. S6. Bottom, histogram showing the distribution of total acceleration events per gene, with the top 5% highlighted in dark blue. (**B**) TF genes are connected to more PREs than non-TF genes. Blue box, inter-quartile range with median indicated; whiskers, 90% data interval. (**C**) TF genes have more acceleration events in their total collection of PREs than non-TF genes. (**D**) PREs of TF genes are more frequently accelerated than non-TF genes, although only slightly so. (**E**) Signaling pathways tend to be connected to large numbers of PREs. Histogram showing a resampling distribution for the numbers of PREs per gene for 50 random genes (Methods). Black bars show the average number of PREs per gene for selected signaling pathways. Dotted line, significance cutoff (*P*_BH_ < 0.05) for resampling test. (**F**) Among those signaling pathways, Notch signaling stood out as showing a high number of total acceleration events per gene (*P*_BH_ = 0.0090; statistics for other pathways are reported in Tab. S6).

To further explore the finding that genes involved in development and gene regulation are enriched for acceleration in their associated PREs, we considered transcription factors (TFs) specifically because of their central role in gene regulation overall and because many TFs are involved in developmental processes. Developmental TFs have been shown to often possess a modular gene regulatory architecture, in which a number of enhancers each drive expression in particular developmental domains [Levine 2010]. Through intersecting our dataset with a hand-curated list of TFs [Lambert et al. 2018], we found that TF genes were connected to a mean of 39.5 PREs, while non-TF genes were connected to 17.5 (Mann-Whitney *U* = 6.0 · 10^6^, *P* = 1.9 · 10^-44^; Fig. 4B). This indicates that TFs in general have more complex *cis-*regulatory architectures, providing ample opportunity for sequence evolution. Indeed, the total number of acceleration events over all PREs and all tested branches was more than twice as high for TFs than for non-TFs (mean number of acceleration events per gene 14.0 vs 6.6, Mann-Whitney *U* = 6.0 · 10^6^, *P* = 1.2 · 10^-44^; Fig. 4C). We also found that PREs that are connected to TF genes were accelerated more frequently than those of non-TF genes, although the difference was small (mean frequency of acceleration 0.60% vs. 0.57%; Mann-Whitney *U* = 7.4 · 10^6^, *P* = 4.8 · 10^-5^; Fig. 4D). These findings are in line with the enrichment results above: Gene regulation is not enriched among genes connected to the most frequently accelerated PREs, and the PREs of TFs are barely more frequently accelerated than PREs of non-TFs. In contrast, gene regulation is enriched among genes having large numbers of PREs and total acceleration events, and TF genes show high numbers of PREs and total acceleration events. Importantly, we found that TFs are affected by a large amount of acceleration over their large number of PREs combined (Discussion).

Building on this series of findings, we were interested to identify specific developmental signaling pathways that were particularly enriched among genes with high PRE connectivity or acceleration within their PREs. We used the KEGG database to select the core genes (as defined by KEGG) from major signaling pathways [Kanehisa et al. 2016] and compared their numbers of PREs with those of resampled gene sets of 50 random genes (Methods). Hedgehog, hippo, MapK, mTOR, Notch, TGF-β, and Wnt signaling all showed larger than average PRE numbers per gene (resampling *P*_BH_ < 0.05; Fig. 4E, Tab. S6), indicating that the regulatory networks of their genes tend to be rather complex. Among those, only Notch signaling showed a higher-than-average total number of acceleration events per gene (18.9 vs genome-wide average, 8.13; resampling *P*_BH_ = 0.009; Fig. 4F, Tab. S6). The fact that only Notch signaling showed an enrichment for acceleration within the PREs of its genes suggests that more extensive gene regulatory innovation may have occurred in Notch signaling than in other signaling pathways.

Further exploring acceleration in Notch signaling-associated PREs, we found that PREs involved with Notch signaling were accelerated on 89 of the 100 branches that we studied (Fig. 5A). As with the overall dataset, long and deep branches showed a lot of acceleration associated with Notch signaling as well: Boreoeutheria had 83 PREs accelerated that were associated with 19 different core Notch signaling genes. Atlantogenata and Sciuromorpha had PREs accelerated that contact 20 different Notch signaling genes each (56 and 40 PREs, respectively). Many individual Notch signaling genes had PREs accelerated on multiple branches (Fig. 5B): Of the 47 Notch signaling genes for which we had data, only two (*APH1A* and *HDAC1*) showed no acceleration at all, three were accelerated on one branch only (*ADAM17*, Australidelphia; *HDAC2*, Caprini; and *KAT2B*, Caprinae), and the remaining 43 were accelerated on multiple branches each (Tab. S7). For instance, *JAG1* had PREs accelerated on 30 different branches, *HES1* had PREs accelerated on 27 different branches, and *NOTCH1* had PREs accelerated on 26 different branches (Fig. 5B). Some individual PREs were accelerated on several different branches (33.8%; Tab. S7). As one example, of the 64 PREs contacting *NOTCH1*, 36 were accelerated, and of those, 23 were accelerated on multiple branches. Overall, these results paint a picture of pervasive sequence acceleration affecting genes of the core Notch signaling pathway.

**Fig. 5.**
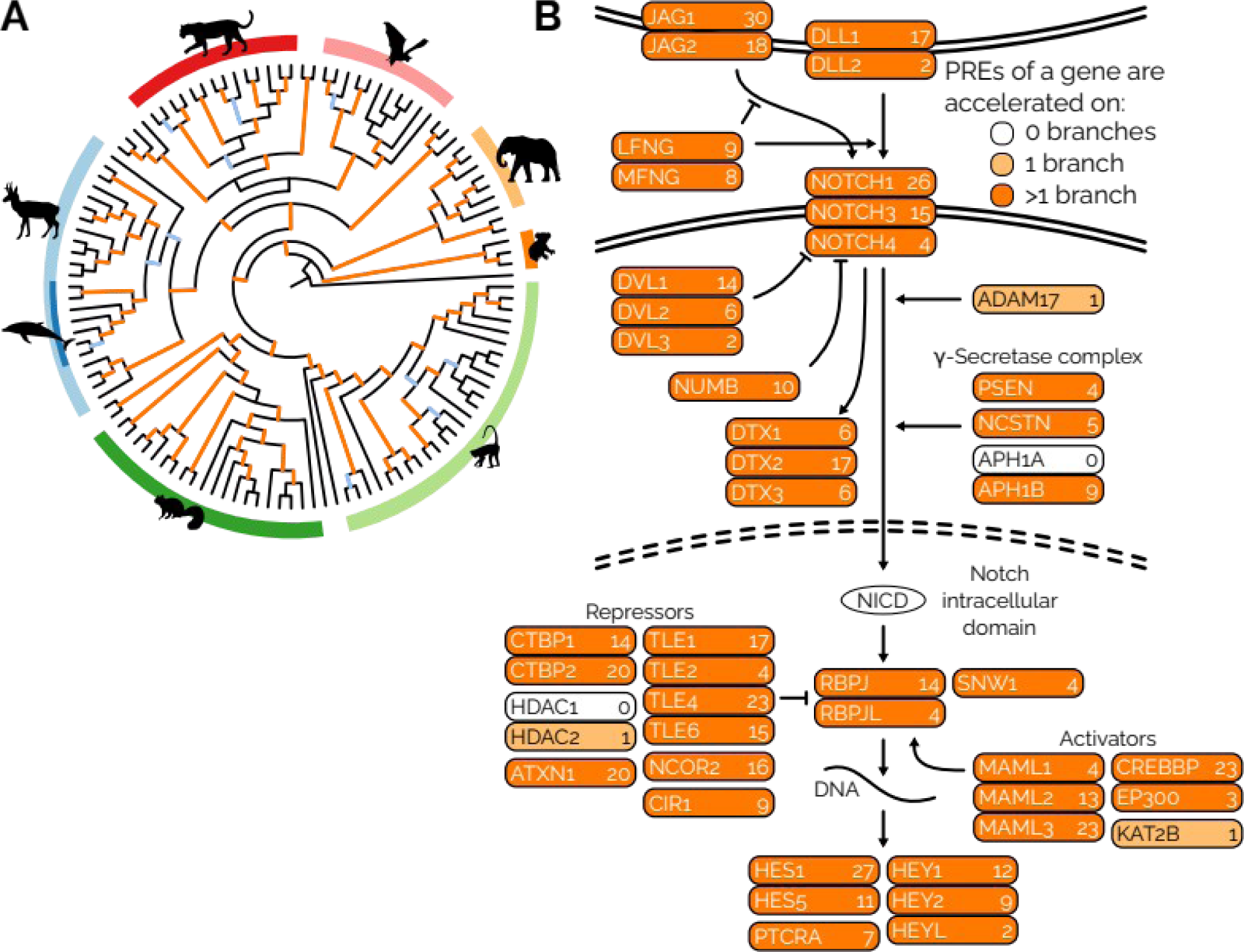
**Acceleration within PREs widely affects the Notch signaling pathway**. (**A**) Tree of life of the whole genome alignment dataset with branches accelerated in PREs that are associated with Notch signaling highlighted in orange. Of the 100 internal branches tested, only 11 (light blue) showed no acceleration in PREs associated with Notch signaling genes. Clade block colors as in Fig. 1. (**B**) The core Notch signaling pathway according to the KEGG database [Kanehisa et al. 2016] with proteins highlighted according to the amount of acceleration observed in their associated PREs. Of the 47 core Notch signaling genes for which we had data, 45 showed acceleration within their PREs and 43 showed acceleration in more than branch. Numbers behind gene names show the number of branches on which PREs contacting the gene were accelerated (Tab. S7).

### Sequence acceleration leading to a gain of an enhancer activity domain

To investigate specific biological impacts of sequence acceleration in a PRE, we designed a lacZ reporter assay to test reconstructed PRE sequences for the ancestral state before, and the derived state after an acceleration event. Building on our finding above that the PREs of genes in the Notch signaling pathway were enriched for sequence acceleration, we focused on PREs associated with Notch signaling genes. We overlapped our set of accelerated PRE sequences with sequences that have been tested earlier in lacZ assays available in the VISTA enhancer browser [Visel et al. 2007]. We found a strong candidate VISTA element in hs1344 (chr3:193,943,028-193,944,689, hg38; Fig. 6A) overlapping four accelerated PREs that are predicted to interact with *HES1*, a core target gene of the Notch signaling cascade [Jarriault et al. 1995, Ohtsuka et al. 1999]. Considering the quality of the alignment and the position of the accelerated branch in the tree, we decided to test an acceleration event that occurred on the branch leading to Pecora, horn-bearing, even-toed ungulates, including eleven species of cows, sheep, and deer in the whole-genome alignment (Fig. 6B). We reconstructed two ancestral sequence versions of the VISTA element (Methods): the ancestral sequence of all pecorans as the *derived* sequence after the acceleration event, and the ancestral sequence of pecorans together with cetaceans, the closest outgroup of Pecora in our alignment, as the *ancestral* sequence prior to the acceleration event. We synthesized both sequences and tested them in an embryonic mouse lacZ reporter assay (Methods).

**Fig. 6.**
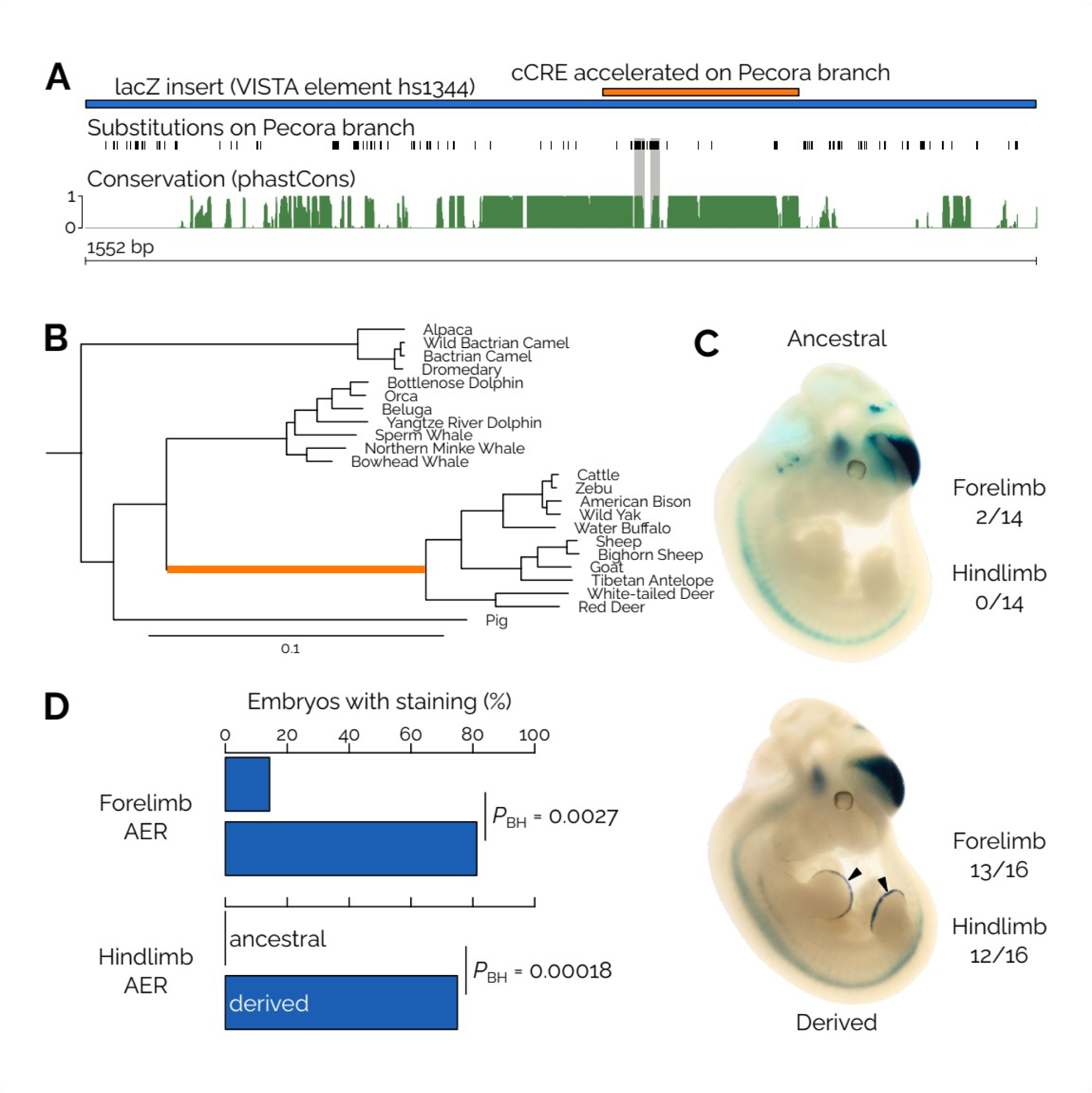
An acceleration event leading to a gain of an enhancer activity domain. (**A**) Overview over the locus. The sequence (orange) found to be accelerated on the branch leading to Pecora sits within a region of conserved sequence. The bulk of the substitutions (black tick marks) that contribute the signal underlying the sequence acceleration event fall into two small parts of conserved sequence surrounded by less conserved sequence (gray bars connecting substitutions and phastCons track, green). Most of the other substitutions that exist within the tested lacZ insert (blue) fall into unconserved sequence space. (**B**) Subtree containing the Artiodactyla species in the alignment with the accelerated branch leading to Pecora highlighted in orange. Branch length scale, genome-wide substitutions per site. (**C**) Expression patterns from the ancestral and derived reconstructed insert sequences driving expression of a lacZ reporter gene in day 11.5 mouse embryos. Arrowheads point to the gained expression domains in the fore- and hindlimb AER in the embryo injected with the derived allele sequence. Numbers on the right indicate the numbers of lacZ positive embryos with staining in the fore- or hindlimb AER. (**D**) Percentage of embryos with lacZ staining in fore- and hindlimb AER for the two alleles. Statistics for additional frequently stained expression domains are shown in Fig. S5.

The experiment resulted in strong and consistent lacZ staining in several domains in both the ancestral and the derived allele (Fig. S5). Interestingly, we found highly reproducible staining in the apical ectodermal ridge (AER) in both forelimbs (13/16 embryos with forelimb staining among lacZ-positive embryos) and hindlimbs (12/16) in embryos carrying the derived allele but not in those carrying the ancestral allele (2/14 and 0/14 in fore- and hindlimb, respectively; Fig. 6C, D). The finding of a new expression domain of a conserved *HES1* enhancer in the AER of the developing limb suggests that the genetic changes of the derived allele sequence modifies gene regulation during distal limb development in horned ungulates. HES1 has been shown to be involved in developmental gene regulation in the AER, and its genetic manipulation affects digit number [Sharma et al. 2021]. Early pecoran fossils show limbs with 4–5 digits [Vislobokova & Trofimov 2002], while all modern members of this group have limbs with two digits, posing the possibility that this *HES1* enhancer may be involved in the evolution of limb development in this group of mammals.

## Discussion

In this study we explored the evolution of gene regulatory elements throughout the mammalian phylogeny by searching for significantly increased substitution rates in putative regulatory element sequences (PREs) on individual branches of the tree. We found this footprint of regulatory evolution to be pervasive across the whole dataset. Genes that saw the highest amount of acceleration among their PREs were enriched for functions in gene regulation and development, and included deeply conserved developmental signaling pathway genes. This matches theory predicting that changes in gene regulation can lead to evolutionary change when protein-coding changes are impeded by pleiotropic constraints.

Theoretical considerations on the evolvability of genetic loci suggested that evolutionary innovation is most likely to occur at loci that minimize pleiotropic effects and at the same time maximize the phenotypic effects [Gompel & Prud’homme 2009, Stern 2013]. Evolutionary change in conserved developmental pathways may have strong phenotypic effects, but detrimental pleiotropic effects if the protein-coding sequence or the promoter sequence is changed, which would affect the activity of the gene everywhere it is expressed. Modularity has been proposed as a way of avoiding strong pleiotropic effects [Wagner et al. 2007]. One way how modularity can be realized regarding the activity of a gene is through a complex network of regulatory elements that each regulate gene activity in distinct contexts. We found that transcription factors, and genes involved in gene regulation and development in general, are enriched for having extensive collections of gene regulatory elements, and for experiencing abundant acceleration over all their PREs and over all branches combined. On any single branch adaptation may involve a variety of diverse biological traits and mechanisms. But overall, evolutionary adaptation in mammals commonly affects modification of gene regulation in conserved, pleiotropic genes involved in gene regulation and development.

Sequence acceleration has previously been found to affect genes with functions related to gene regulation and development in studies investigating single branches in the mammalian tree of life [Pollard et al. 2006, Holloway et al. 2016, Bi et al. 2023, Zhuang et al. 2023]. When we inspected acceleration on a single branch, this pattern was not always apparent, but it was prominent in our analysis across the entire mammalian tree. This illustrates the utility of investigating acceleration at this scale. Focusing on a single branch or a single trait is not sufficient to provide general insight into which genes and traits are frequently experiencing recurrent evolutionary change. Assessing the evolution of regulatory elements across an entire, large clade, and without biases for certain traits or groups of genes allowed us to draw generalizable conclusions on the types of genes that are predominantly affected by regulatory evolution. We also took advantage of analyzing acceleration as a relative frequency across lineages instead of inspecting lists of accelerated regions in isolation. This approach provided a baseline for the amount of acceleration observed, and gives a more nuanced picture of which types of PREs and associated genes experience more acceleration than others.

Genes involved in Notch signaling were particularly enriched for having experienced a large number of acceleration events, and several core Notch signaling genes showed acceleration events in well over a dozen branches. Notch signaling assumes a central role in animal development and has connections to many other developmental signaling pathways, a fact that may make it an important substrate for evolution [Guruharsha et al. 2012, Nian & Hou 2022]. When we tested the regulatory effect of an acceleration event in a PRE interacting with *HES1* in a lacZ reporter assay in the developing mouse embryo, we found that the derived allele resulting from the acceleration event drove strong reporter activity in the apical ectodermal ridge (AER) of both developing limbs. The ancestral allele prior to the acceleration event was not active in the hind- and only sporadically active in the forelimb. The AER is an important signaling center in the developing limb [Fernandez-Teran & Ros 2008]. HES1 has been reported previously to regulate target gene expression in the AER with genetic perturbation of HES1 signaling affecting digit number [Sharma et al. 2021]. This is of potential biological relevance, as all modern pecorans show reduced digit numbers in their limbs, while early pecoran fossils show limbs that more closely resemble the general mammalian type with five digits [Vislobokova & Trofimov 2002]. This poses the intriguing hypothesis that this acceleration event affecting developmental gene expression in the AER on the branch leading to Pecora could affect digit number and may have played an important role in the adaptive evolution of locomotion in this group of mammals.

We further provide evidence that sequence acceleration affects gene regulatory function more generally. We found that acceleration events tended to coincide with increased rates of transcription factor binding site (TFBS) predictions. While *in silico* predictions of TFBSs do not always reliably reflect specific *in vivo* binding events, the pattern we found over our entire dataset can be expected to include a sufficient amount of true binding events. We also found that acceleration events tended to coincide with turnover in chromatin accessibility predicted using machine learning [Kaplow et al. 2023]. The overall pattern we found was somewhat weak, which we believe is mostly due to the fact that the number of loci overlapping between our dataset of accelerated PREs and the analyzed open chromatin dataset is small. However, sequence differences need not necessarily affect chromatin accessibility. A regulatory element sequence may remain accessible despite changes in regulatory activity [Oruba et al. 2020]. An acceleration event may also affect regulatory activity in a different cell type than the one that was assayed for chromatin accessibility. Overall, our findings indicate that sequence acceleration as identified in this study can modify regulatory element activity through sequence content turnover.

To identify factors that influence a PRE’s propensity to be accelerated, we interrogated a range of putative explanatory correlates. Branch length, i.e., evolutionary divergence, was correlated with acceleration frequency. This reflects that more time provides more opportunity for acceleration events to occur. Evolutionary sequence conservation also affected the levels of acceleration that we observed, in several ways. We found lower acceleration frequencies and thus higher constraint in cases where 1) the PRE itself was constrained, as measured by phastCons; 2) the contacted gene was constrained, indicated by gnomAD data; 3) the amount of regulatory space was limited, in cases where genes were contacted by small numbers of regulatory elements. These findings show that sequence acceleration is affected by known evolutionary constraints. It is interesting to note the central role assumed by a PRE’s target gene in determining some of the constraints, both through the gene’s own level of constraint and through the extent of the gene’s regulatory space.

There are some biases and caveats that apply to our study. We ascertained our set of PREs based on evolutionary conservation. PREs were defined based on phastCons elements generated from the 120- mammal alignment, or based on conserved biochemical activity between human and mouse. This excludes any regulatory elements that are restricted to a small number of species, or to genome regions that cannot be aligned to the human reference genome. Hence, our dataset is biased towards conserved regulatory elements and enriched for such elements that regulate the activity of conserved genes. It is noteworthy that even within this dataset, we found an enrichment for accelerated sequence evolution affecting conserved, developmental signaling genes. However, studying younger gene regulatory elements, or elements that are not as conserved and that turn over more frequently may produce different results than reported here. While a more inclusive approach may paint a more complete picture of evolutionary change in gene regulatory elements, our approach of focusing on deeply conserved elements provides nuanced insight into how evolution affects some of the most ancient and fundamental mammalian genes and pathways.

## Methods

### Data collection

The analyses in this paper were performed on a previously published whole-genome alignment of 120 mammal species [Hecker & Hiller 2020]. To collect a set of regulatory sequences, we selected two types of putative regulatory elements (PREs) for this study, phastCons elements (pCEs) and ENCODE cCREs. pCEs were downloaded from the Hiller lab website (https://bds.mpi-cbg.de/hillerlab/120MammalAlignment/). pCEs less than 10 bp apart were merged before those shorter than 51 bp were excluded to select for sequences long enough to produce acceleration test results with sufficient statistical support, but there was no such limit at the upper end (maximum pCE length 1886 bp; average length, 134.2 bp). To select ENCODE cCREs, we downloaded the full human (hg38) cCRE set that was available in October 2020. We also downloaded ChIP-seq-derived chromatin activity data from mouse (ENCODE cCREs and ENCODE developmental ChIP-seq derived ChromHMM tracts [Gorkin et al. 2020] identified as enhancers or promoters), lifted it over to hg38 (--minMatch=0.8) and kept those human cCREs that overlapped mouse activity data. ENCODE cCREs range in size between 150 and 350 bp, with a mean length of 271.5 bp. We filtered both types of PREs against promoters (1 kb upstream of transcription start sites), exons and pseudogenes (hg38 Gencode v.40), and repeats (UCSC genomicSuperDups, Simple Repeats and RepeatMasker annotations and the hg38–hg38 self alignment chain). The final PRE dataset contained 319,292 pCEs and 115,014 ENCODE cCREs.

To determine enrichment of overlap of cCREs with pCEs, we used BEDTools shuffle to generate 10,000 random sets of sequences of the same length as our cCREs within the filtered genome. Between 9.9% and 10.8% of reshuffled cCREs overlapped pCEs, and none reached the amount of overlap of the original set (43.8%).

### Testing for sequence acceleration and branch-specific quality filters

We selected 100 internal branches in the 120 mammal alignment tree. To be able to distinguish segregating genetic variation from lineage-specific variants, at least two genomes need to be present on either side of the tested branch, which is why we were restricted to the use of internal branches. We did not test all internal branches, because many are rather short, with the consequence that there is limited genetic information separating a specific branch from the rest of the tree. This is particularly problematic for short but old branching events, where additional substitutions may have overwritten an earlier pattern. To arrive at the final set of branches, we manually inspected the tree, focusing on a branch’s length and its position in the tree, and included as many branches as possible that we deemed long and insightful enough. We then used phyloP to test for acceleration (-o ACC) using its likelihood ratio test method (-m LRT) on each branch (-B) [Pollard et al. 2010, Holloway et al. 2016].

To exclude cases where a pattern of acceleration is likely due to GC-biased gene conversion, we ran phastBias per branch with the option to output GC-biased gene conversion tracts [Capra et al. 2013]. Tracts that were up to 1 kb apart were merged and PRE acceleration results were filtered from the dataset if they overlapped gene conversion tracts on their respective branch.

Finally, we filtered out PREs on a branch-wise basis depending on their clade-specific alignment quality. For every clade, we required that there was at least one species in each of the two branches immediately underneath the focal branch that had at least 80% and at most 120% of bases present in the alignment, relative to the reference sequence (hg38). We found that this excluded low quality alignments well while retaining alignments with putatively true patterns of acceleration even in PREs with lower overall levels of sequence conservation.

### Connecting PREs to potential target genes and gene ontologies

We connected PREs to their potential target genes using two strategies. First, we used whole-genome chromosome conformation capture-type datasets from developmental primary samples or cell models. Second, we associated PREs with nearby genes if a PRE overlapped the range 1–5 kb upstream of a gene’s transcription start site. Chromosome conformation capture-type data was combined as follows: Human neural progenitor cell pcHiC data [Jung et al. 2019] was provided as a list of gene names interacting with genomic intervals in hg19. Data was lifted to hg38 and gene names were translated to ENSEMBL gene identifiers using a dictionary from biomaRt. Human PLAC-seq data from radial glia, intermediate progenitor cells, excitatory neurons, and interneurons [Song et al. 2020] was provided as interacting genomic intervals in hg38. Both intervals were separately BED-intersected with gene promoters (5 kB upstream to 1 kB downstream of transcription start sites) from Gencode v40. Human HiC data for primary ventricular zone and cortical plate [Won et al. 2016] was provided as genomic intervals interacting with ENSEMBL genes in hg19, which were lifted to hg38. Human HiC data from neural progenitor cells, neurons, and astrocytes [Rajarajan et al. 2018] was provided as interacting genomic intervals in hg19. Both intervals were separately BED intersected with Gencode v40 promoters (in hg19) and lifted to hg38. Human fetal heart DNase HiC data [Bertero et al. 2019] was provided as a contact matrix in hg38. We ran Arrowhead (Juicer Tools v 1.22.01) to find interacting domains and BED intersected both interacting intervals separately with Gencode v40 gene promoters. Mouse postnatal day 28 cortex HiC data [Du et al. 2017] was provided as a contact matrix in mm9. After running Arrowhead, data was lifted to mm39, and interacting intervals were separately BED intersected with Gencode vM29. Mouse ENSEMBL identifiers were translated to human using a dictionary from biomaRt. Once all data were in the form of human ENSEMBL gene identifiers interacting with genomic intervals, these intervals were BED intersected with our PRE dataset.

We annotated genes as transcription factors according to the classification in [Lambert et al. 2018] (with one addition [Grand et al. 2021]) and as non-transcription factors all genes that were not evaluated in that study.

The human orthologs from core pathway genes of ten major signaling pathways (Hedgehog, Hippo, JAK-STAT, MAPK, mTOR, NF-κ B, Notch, Ras, TGF-β, Wnt) were retrieved from the KEGG pathway database [Kanehisa et al. 2016]. We compared genes from these pathways with those of resampled sets of 50 random genes. We chose 50, because the smallest pathways comprised 47 and 50 genes, respectively. Choosing a small number leads to a large variance in the resampling distribution which leads to more conservative test results. We compared the numbers of associated PREs, the relative acceleration frequency, and the number of acceleration events per gene between pathway and randomized gene sets.

### Data analysis

Statistical analyses were performed using R version 4.2.0. Gene Ontology and Reactome annotations were downloaded using the BioConductor package biomaRt (v. 2.52.0). Enrichment testing was done using hypergeometric tests. GOslim annotations for all significantly enriched biological process GO terms in the per-branch analysis were retrieved from EBI’s QuickGO. GOslim category counts were log_2_ transformed and normalized to the total number of GO term counts per branch for visualization.

We mapped transcription factor binding site (TFBS) archetypes [Vierstra et al. 2020], containing position weight matrices from three sources [Jolma et al. 2013, Khan et al. 2018, Kulakovskiy et al. 2018] to all genome sequences in the alignment. We intersected TFBS predictions with our PRE dataset using BED intersect (BEDTools v2.30.0), requiring that the full TFBS prediction overlapped a PRE (-f 1). We then calculated the asymmetric binary distance of the TFBS sets in any PRE between all species within the ingroup and the outgroup of a focal branch. For this branch we then compared the mean distance for all ingroup and outgroup species with the difference of means of the ingroup and the outgroup for cases where said branch was accelerated versus cases where the branch was not accelerated.

Multi-species open chromatin region (OCR) predictions for three different samples were downloaded from http://daphne.compbio.cs.cmu.edu/files/ikaplow/TACITSupplement/PredictionMatrices/ [Kaplow et al. 2023]. Predictions were based on data from on four (motor cortex), five (liver), and two species (PV^+^ interneurons), respectively. The species for which predictions had been generated did not fully overlap with our dataset such that we were able to compare data for to 83 (motor cortex/liver) or 36 branches (PV^+^ interneurons) per sample. We intersected OCRs with our PRE dataset using BED intersect, requiring that at least 50% of our PREs were overlapping (-f 0.5). We did not require a minimum overlap in the inverse direction (-F), because OCRs were much larger than most PREs, such that in most cases, the length of a PRE is <50% the length of an OCR. OCRs are assigned a predicted value for their accessibility ranging from 0 (closed) to 1 (open). For each analyzed branch, we formed the absolute difference between the mean of this value of the ingroup and of the outgroup (Fig. 3C). We then tested if this value differed between regions where the branch was accelerated and those where the branch was not accelerated. Many of the tests were insignificant, because the number of OCRs overlapping PREs was small, and because acceleration was not very common on many branches.

### LacZ reporter assay

To select a sequence to test in a lacZ assay, we intersected accelerated PREs that were predicted to interact with genes in the core Notch signaling pathway with all VISTA elements [Visel et al. 2007], *bona fide* enhancers with known expression domains. After inspecting the expression patterns of the resulting VISTA elements, we decided to target hs1344 (chr3:193,943,028-193,944,689, hg38), which showed strong and consistent activity in a lateral forebrain domain. We extracted sequences orthologous to hs1344 from each genome sequence in the clade Artiodactyla, aligned them using Clustal Omega (www.ebi.ac.uk/Tools/msa/clustalo/), and corrected alignments manually.

We then reconstructed ancestral sequences at our two nodes of interest (before and after the acceleration event on the Pecora branch) using baseml (PAML v. 4.9j [Yang 2007]; options verbose=2, runmode=0 [user-provided tree], model=7 [general time reversible, REV], clock=2 [local clock, with differing rates at the Pecora branch and in the rest of the tree], fix_kappa=0 [estimate kappa], fix_alpha=0 [estimate alpha], RateAncestor=1 [estimate ancestral states]). Baseml can either discard or include all ambiguous characters found in the alignment for estimation. In the case of insertions and deletions, this means that either all sites that contain a deletion in any of the sequences of the alignment are removed from the output, or that all insertions are included in the output, in each case including species-specific deletions and insertions that are irrelevant for our nodes of interest. Hence, we did not remove ambiguous sites (option cleandata=0) and cleaned the reconstructed sequences from superfluously inserted sites by hand.

We also removed 183 bases from the 3’ end of the insert sequence because they contained a low complexity sequence that was difficult to synthesize. The resulting *ancestral* (before the acceleration event; 1525 bases) and *derived* (after the acceleration event; 1526 bases) sequences (Tab. S8) were then synthesized as gBlocks HiFi Gene Fragments by IDT. Synthesized sequences were cloned into an *Hsp68-lacZ* reporter vector through Gibson assembly (GeneArt #A46624) using the primers listed in Tab. S8. Generation of transgenic mice and staining of day 11.5 embryos were carried out as previously described [Pennacchio et al. 2006]. Stained mouse embryos were photographed from four sides (left, right, ventral, dorsal) and scored independently by three blinded scorers.

### Data availability

Code is deposited at GitHub: https://github.com/severinEvo/Uebbing_et_al_acceleration. All tested and significantly accelerated PREs are available as BED file at Zenodo: <u>10.5281/zenodo.10600206</u>.

## Acknowledgments

This work was supported by National Institutes of Health awards R01 HD102030 (to J.P.N.) and F32 HD108935 (to M.B.). This research program and related results were also made possible by the support of the NOMIS foundation (to J.P.N.). We thank all Noonan lab members for their helpful discussions and L. Pennacchio for providing a list of VISTA elements.

## Contributions

S.U. and J.P.N. conceived of and designed the study. S.U. performed computational analyses. S.U. performed experiments with help from A.A.K., M.B., and Y.J. S.B., X.X., and T.N. injected mouse embryos. S.U. wrote the text with input from all authors.

## Supplementary Figures

**Fig. S1.**
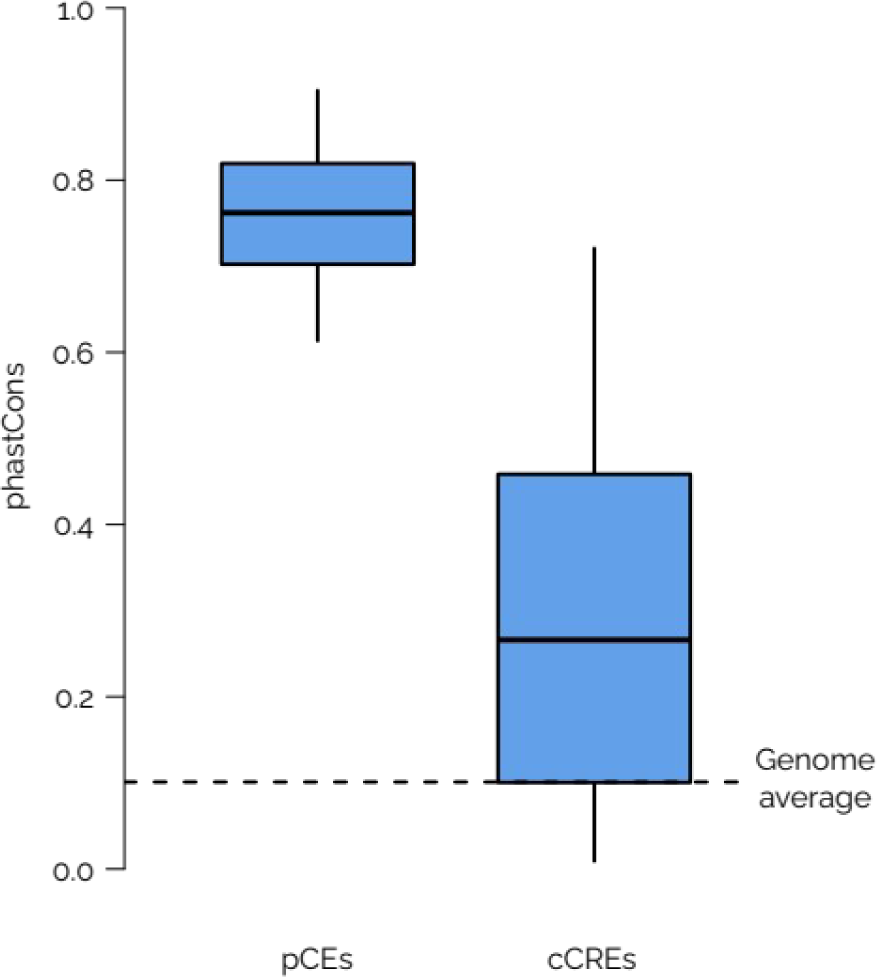
Both types of PREs show elevated phastCons scores. PhastCons scores of pCEs and ENCODE cCREs compared to the average genomic phastCons score (dotted line) exclusive of genes and repeats. pCEs are defined using phastCons, but even ENCODE cCREs show higher phastCons score than the genome average.

**Fig. S2.**
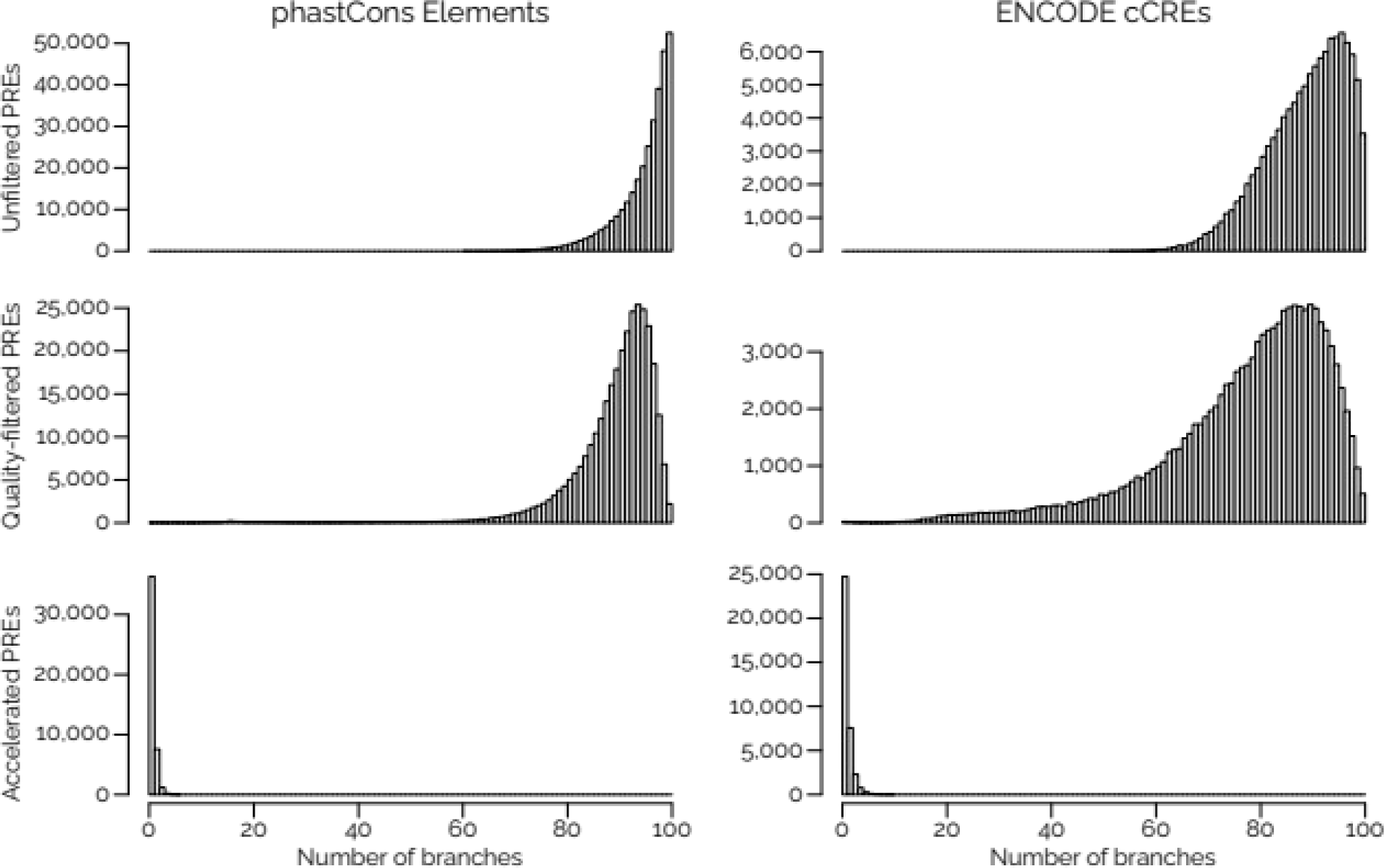
Filtering steps remove few PRE-branch instances from the dataset. Distributions of the numbers of unfiltered, filtered, and accelerated PREs that are present per analyzed branch show that filtering removes individual PRE-by-branch combinations, but most PREs are still present for the majority of branches. In contrast, acceleration signal is only detected in individual branches, indicating that the testing procedure is conservative and not nearly saturated towards the amount of PREs present. Unfiltered PREs may be missing from a branch if the alignment does not contain this sequence in some species due to deletions or assembly gaps.

**Fig. S3.**
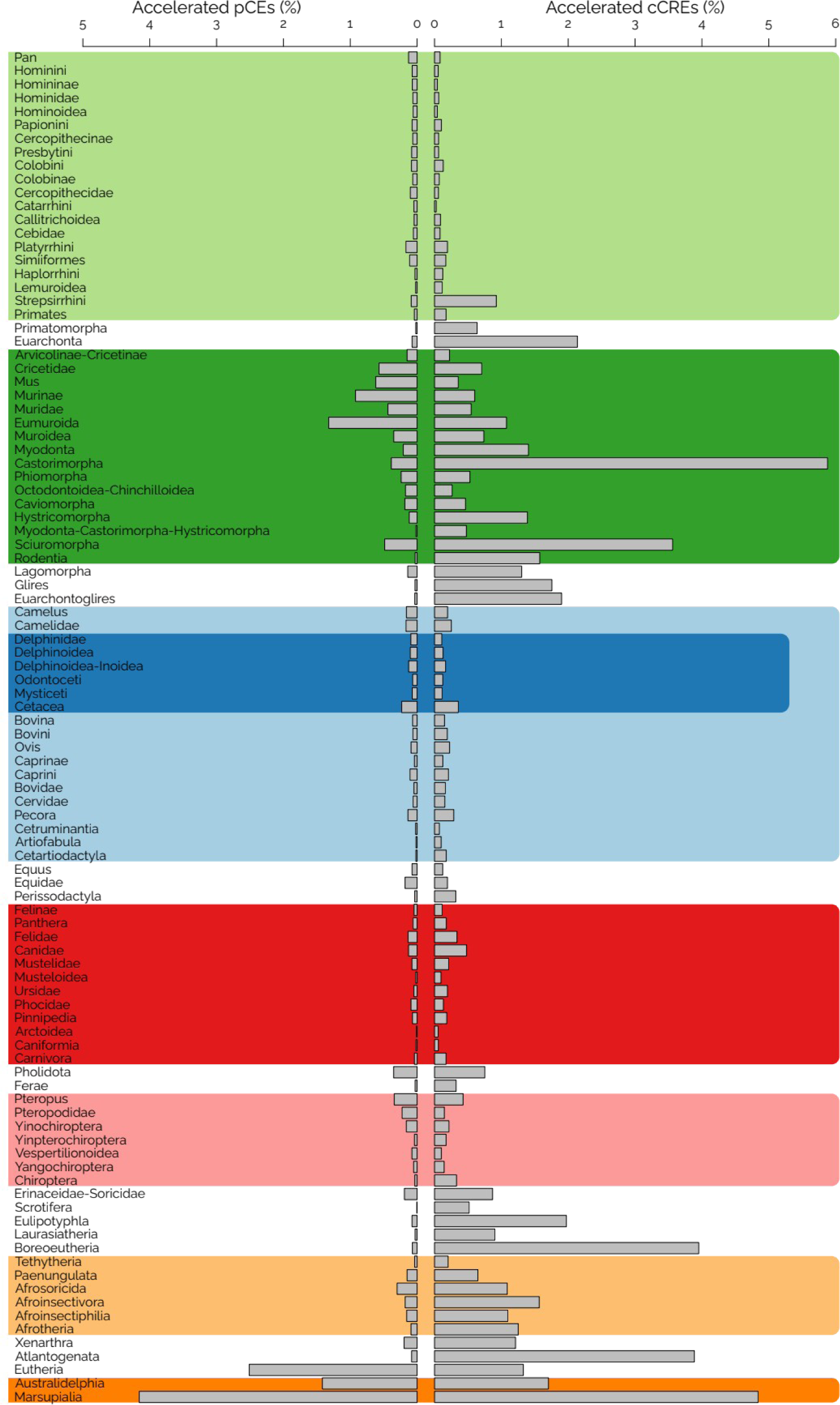
Expanded acceleration results per tested branch and PRE type. See Fig. 1B for details.

**Fig. S4.**
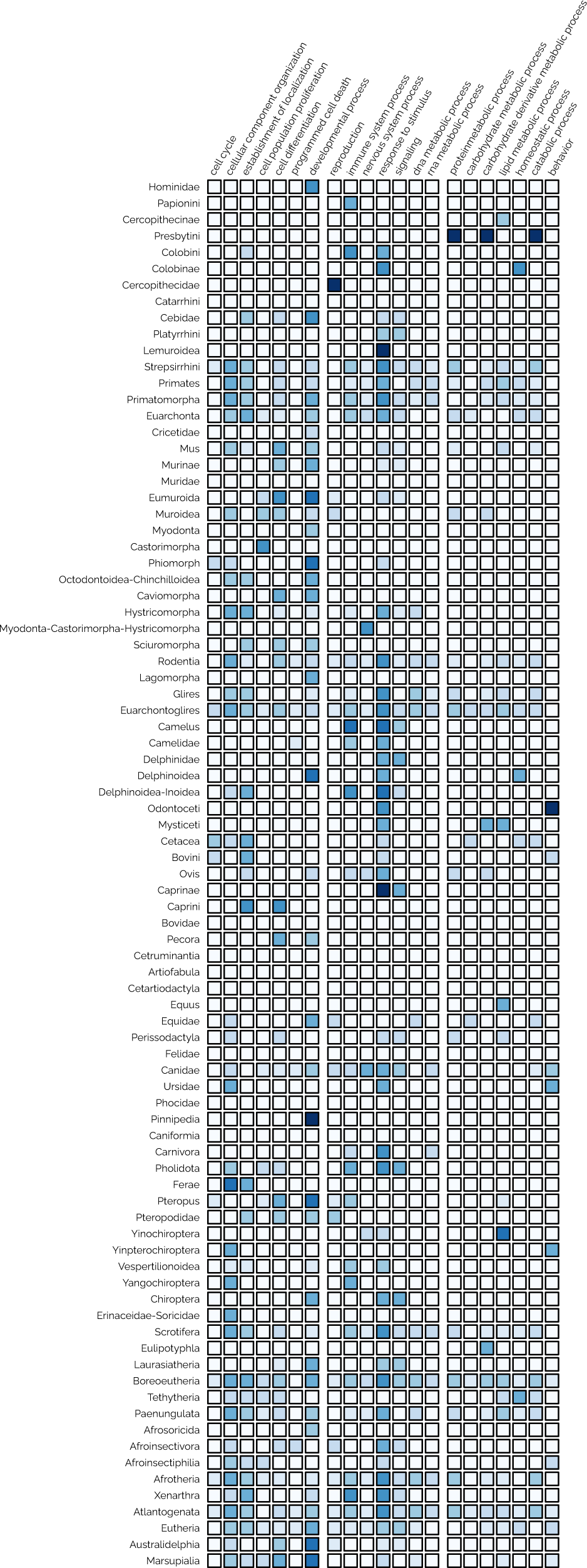
Reduced complexity representation of GO enrichment among accelerated regions per branch. Enriched biological process GO terms were associated (associations from ENSEMBL) with GOslim categories from the Alliance of Genome Resources. Darker blue shading indicates a stronger enrichment with GO terms within this GOslim category. Data in each row is normalized to the total number of enriched GO categories by branch.

**Fig. S5.**
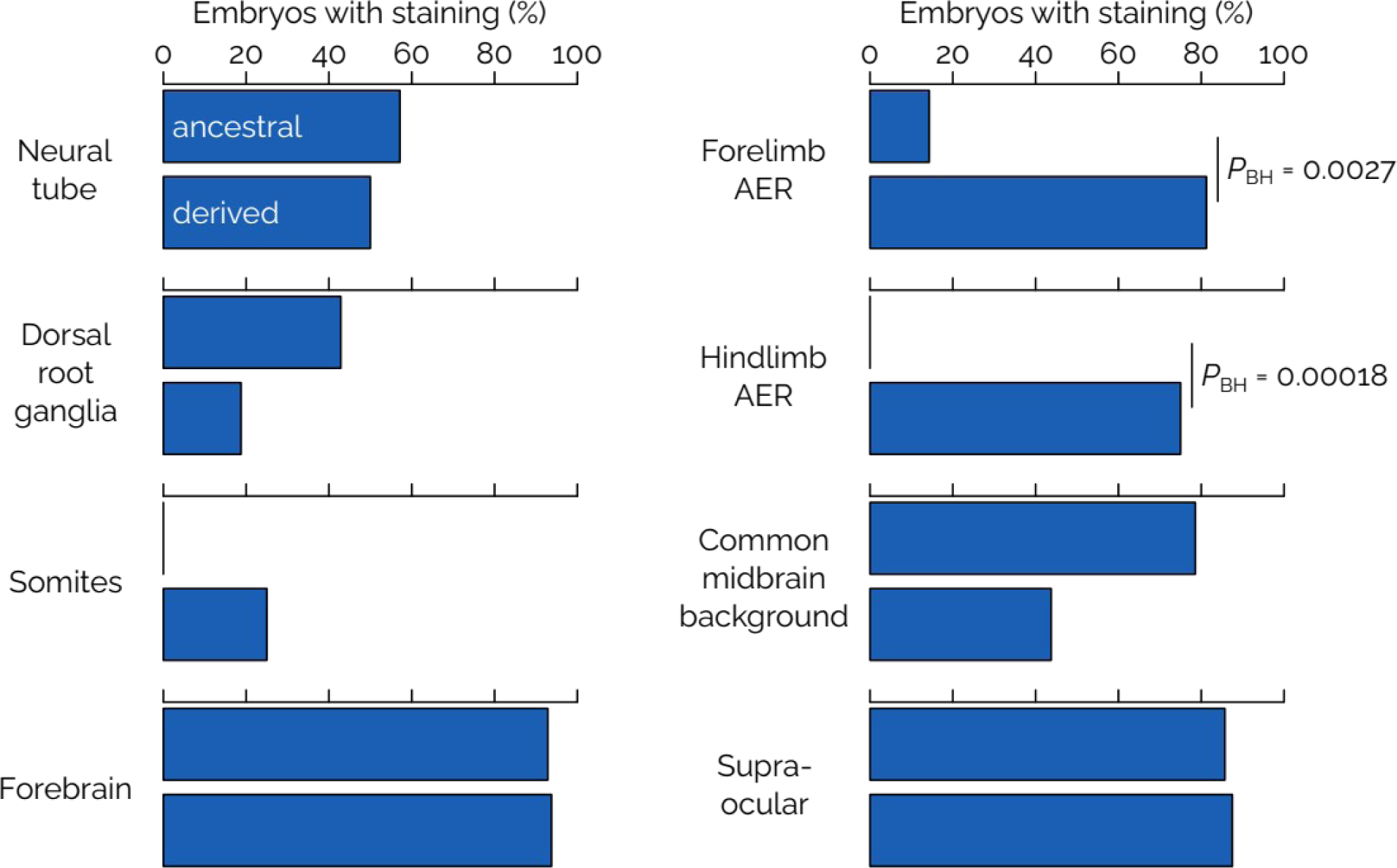
Frequently staining lacZ expression domains. Frequency of lacZ staining for the two alleles in commonly staining expression domains. The ancestral allele is always plotted in the top, the derived allele in the bottom. Total numbers of lacZ positive embryos were n = 14 for the ancestral allele and n = 16 for the derived allele.

